# Hexafluoro slows retinal degeneration and improves visual function in zebrafish models of Usher syndrome 1F

**DOI:** 10.1101/2023.12.29.573664

**Authors:** Jennifer B. Phillips, Siena Kulis, Sara H. Buchner, Eric J. Fox, Jeremy Wegner, Judy Peirce, Maryna V. Ivanchenko, David P. Corey, Jack L. Arbiser, Monte Westerfield

**Affiliations:** Institute of Neuroscience, University of Oregon, Eugene, OR, USA; Department of Neurobiology, Harvard Medical School, Boston, MA, USA; Metroderm/UDP, 875 Johnson Ferry Rd, Atlanta, GA, USA

## Abstract

Usher syndrome is the leading genetic cause of deafblindness, affecting hundreds of thousands of people worldwide. The deafness can be addressed with hearing aids or cochlear implants, but there is currently no treatment for the vision loss, which is due to progressive degeneration of retinal photoreceptors. Studies in animal models of Usher syndrome have shown that photoreceptor degeneration is exacerbated by exposure to bright light, and other studies have shown that light-induced photostress reduces mitochondrial function. We previously synthesized hexafluoro and showed that it is a potent Sirt3 activator that promotes mitochondrial respiration. Here we examined the efficacy of hexafluoro as a potential therapeutic for treatment of vison loss in a zebrafish model of Usher syndrome type 1F, which exhibits early and severe vision defects along with vestibular dysfunction as seen in Usher type 1 pathology. We find that hexafluoro improves visual function, reduces photoreceptor degeneration, and protects the retina against exposure to bright light in this USH1F model.

## INTRODUCTION

Usher syndrome (USH) is the most prevalent cause of hereditary deafblindness, characterized by congenital hearing loss, progressive retinal degeneration, and in some cases balance problems^[23, 30]^. USH is an autosomal recessive disorder with three clinical subtypes, USH1-USH3, distinguished by age of onset and severity of symptoms, with USH1 being the most severe. Six different genes have been associated with USH1, including *PCDH15*, which is responsible for Usher syndrome type 1F (USH1F). *PCDH15* encodes a transmembrane protein with a large extracellular domain that contributes to the tip-links of the stereocilia of mechanosensory hair cells^[18]^. The function of PCDH15 in the retina is less well understood, but studies have shown that it localizes to the inner segment plasma membrane, calyceal processes, and synaptic regions of photoreceptors^[20, 32]^. USH1F is characterized by congenital profound deafness, vestibular deficit, and progressive vision loss that is a form of retinitis pigmentosa^[6]^.

The pathophysiological mechanisms that underlie retinal degeneration in USH are only partially understood. The retina uses large amounts of energy to convert light into electrical signals^[16, 35]^. Multiple stress response mechanisms, such as the unfolded protein response^[2, 27]^ and DNA repair^[33]^, which are involved in preserving retinal homeostasis during light transduction, require energy in the form of ATP. Regulation of the energy balance is thought to be a key factor involved in neuroprotection against both retinal^[17]^ and inner ear^[14]^ diseases because energy decompensation results in a failure of these stress response mechanisms. In support of this notion, previous studies have shown that light-induced photostress reduces mitochondrial respiratory function and ATP levels in photoreceptors, and restoring mitochondrial function promotes neuronal survival and retinal function^[17, 19]^. We have shown that light exposure hastens retinal degeneration in zebrafish USH models^[9]^, and mitochondrial dysfunction has also been implicated in sensory hair cell death^[13]^. Thus, mitochondria may be an important target for USH therapy.

Sirt3 is an NAD-dependent deacetylase localized mainly in mitochondria where it functions as the primary deacetylase. A mass spectroscopic study revealed that nearly 65% of mitochondrial proteins are acetylated, implying an important role of Sirt3 in mitochondria^[12]^. SIRT3 maintains mitochondrial health by deacetylating a wide variety of enzymes involved in metabolism, reduction of reactive oxygen species, apoptosis, and mitochondrial dynamics^[3]^. Thus, compounds that activate Sirt3 are good candidates for preserving photoreceptors and inner ear hair cells in USH. Previously, we synthesized^[5]^ hexafluoro (bis-trifluoromethyl-bis-(4-hydroxy-3-allylphenyl)methane), a potent small molecule activator of Sirt3, and showed that it promotes mitochondrial respiration and saves cardiomyocytes from mitochondrial damage and cell death induced by doxorubicin^[26]^. Hexafluoro is an analog of honokiol, a natural product derived from the bark of Magnolia. Because honokiol is difficult to extract or synthesize, we generated hexafluoro, which is fluorinated to prevent rapid metabolism^[5]^.

In this study, we used gene editing of the *pcdh15b* gene to generate zebrafish models of USH1F. We characterized the hearing, balance, and vision defects of these mutants and then examined the efficacy of hexafluoro treatment to ameliorate the retinal defects. We show that hexafluoro significantly improves visual function and reduces photoreceptor degeneration in these USH1F models, thus identifying hexafluoro as a promising therapeutic for treating vision loss in USH1F patients.

## RESULTS

### Zebrafish *pcdh15b* mutants have balance and inner ear mechanosensory hair cell defects

The R245X change in PCDH15 is considered a founder mutation responsible for approximatively 50-60% of USH1 in the Ashkenazi Jewish population^[6]^. Given the prevalence and severity of the R245X mutation, we generated zebrafish with similar truncating mutations to use as models to develop effective therapies for this group of affected individuals. Because of the previous analysis identifying *pcdh15b* as the zebrafish orthologue critical for pcdh15 protein function in the retina^[28]^, we used CRISPR/Cas9 to generate small deletions in Exon 8 of the zebrafish *pcdh15b* gene that introduce a frameshift and premature termination codon in the region homologous to the sequence encoding R245 in the human *PCDH15* gene. We obtained two deletion alleles from different founders: *pcdh15b*^*b1257*^, a 7bp deletion (c.755-61; p.L252Pfs*6) and *pcdh15b*^*b1277*^ a 4bp deletion (c.756-9; p.T253Pfs*5). Homozygous mutants for each allele were phenotypically indistinguishable. All experiments described in this manuscript used *b1257* mutants due to a greater number of breeding carriers obtained with this allele, and are hereafter referred to as simply ‘*pcdh15b’*.

Based on the morpholino and mutant phenotypes previously documented^[20, 28]^, we anticipated that homozygous *pcdh15b* larvae would exhibit retinal defects but normal hearing and balance behavior. Mutant *pcdh15b* larvae displayed a normal acoustic startle response, responding to tapping on the edge of the Petri dish 100% of the time. However, in contrast to previous studies^[20, 28]^, *pcdh15b* mutants exhibited marked vestibular defects including impaired righting behavior and swimming in twirling, circling patterns, as documented for other zebrafish mutants with defects in sensory hair cell mechanosensation^[21]^. Examination of the inner ear sensory patches revealed disorganized hair bundles with splayed or bent stereocilia (Fig. 1A-E). Bundle morphology was more severely perturbed in the anterior (utricular) macula of *pcdh15b* mutants (Fig. 1C). More than half of the hair bundles observed in the mutant utricles were bent or splayed, compared to fewer than 5% in wild type. Milder disorganization of the bundles was observed in the posterior (saccular) macula (Fig. 1E: 19% in mutants compared to 9% in wild type). We also noted a reduction in the overall number of hair bundles in both the utricule (39.2 ± 2.05 in mutants compared to 48.33 ± 2.42, *P* = 0.07 in wild type) and saccule (60 ± 0.89 in mutants compared to 71.3 ± 1.03, *P* = 0.006 in wild type) in mutants compared to wild type at 5 days postfertilization (dpf), a phenomenon we reported previously in other USH1 fish models^[4, 25]^. Stereocilia disruptions were milder than those observed in *pcdh15a* mutant larvae^[28]^, but illuminated a hitherto unknown role for *pcdh15b* in the zebrafish inner ear. To our knowledge, this is the first example of a zebrafish mutant having a more severe phenotype than a morpholino knockdown. The vestibular phenotype documented in these mutants is distinct from a recently published zebrafish USH1F genetic model^[20]^, which has normal swimming and balance and survives until 20 dpf. In contrast, the balance defects were severe in animals with truncating mutations in exon 8, and these mutations are early larval homozygous lethal presumably due to the impact of poor vestibular function on the ability of mutants to orient toward, capture, and feed on live prey.

**Figure 1.**
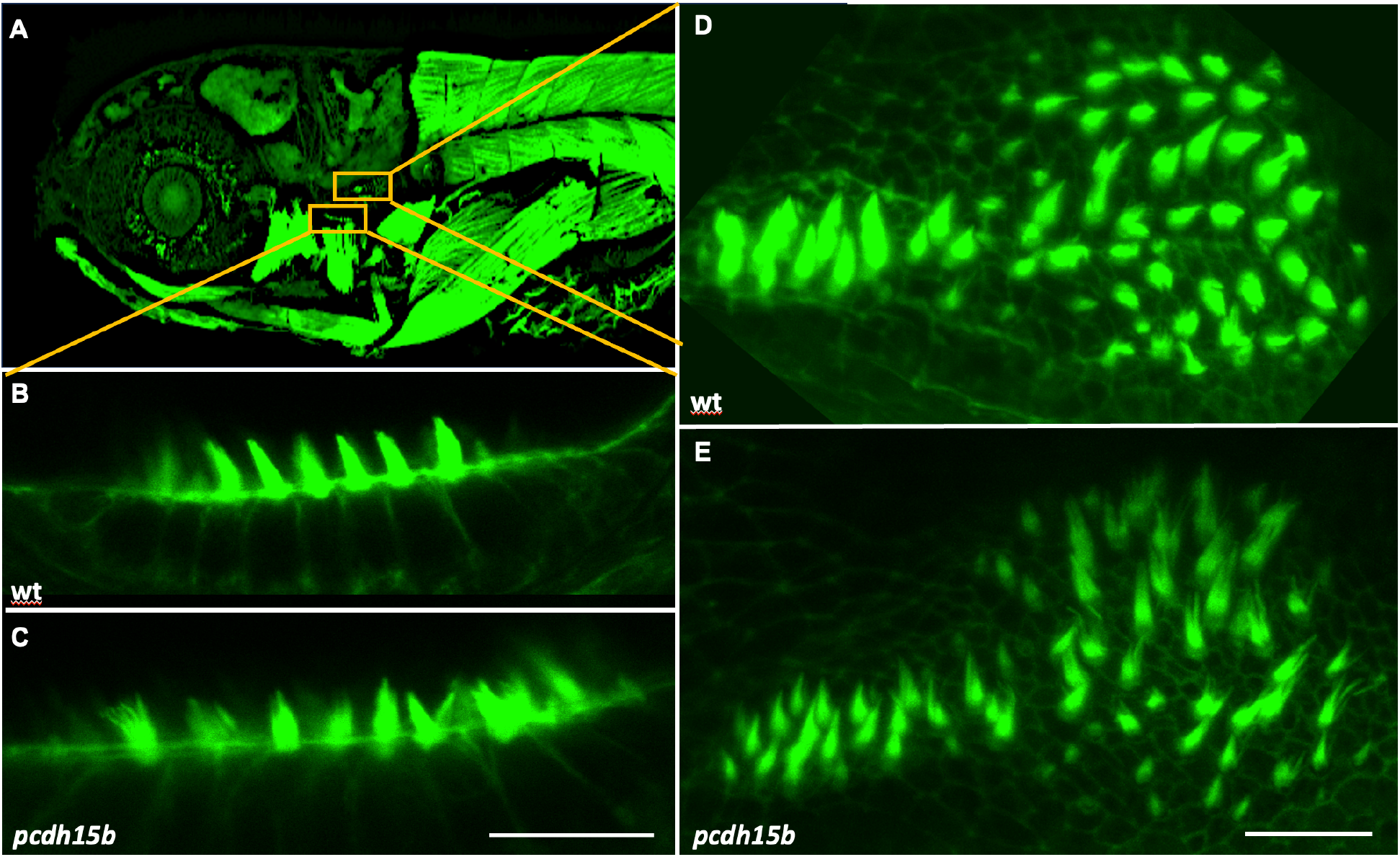
*pcdh15b* mutants have balance defects and disrupted stereocilia. (A) Image of 5 dpf phalloidin labeled larva to provide orientation for panels B-E. (B-E) Hair cells populating the large sensory patches of the inner ear are labeled with fluorescently conjugated phalloidin. (B, D) Macular hair bundles in wild-type zebrafish larvae are organized and tapered at the tips. (C, E) A truncating frameshift mutation in exon 8 of *pcdh15b* results in splayed and bent hair stereocilia throughout the sensory patches of the inner ear. Scale bars: C (10 µm) also applies to B; E (20 µm) also applies to D. *n* = 6 animals for hair cell statistics.

### Zebrafish *pcdh15b* mutants have disrupted photoreceptor outer segments, visual function defects, and photoreceptor degeneration

Histological analysis of *pcdh15b* mutant retinas showed significant disruptions to photoreceptor morphology, consistent with the previous reports from *pcdh15b* mutants^[20]^ and morphants^[28]^. Labeling with an antibody against cone transducin was diminished and abnormally distributed in *pcdh15b* mutants, revealing misshapen outer segments (Fig. 2A, B). Labeling with phalloidin to visualize the calyceal processes showed marked abnormalities in shape, arrangement, and number of these fingerlike processes that ensheath the outer segment (Fig. 2C, D). To investigate these changes more precisely, we performed scanning electron microscopy and transmission electron microscopy on zebrafish retinas (Fig. 2E-H). In *pcdh15b* larvae, the outer segments of photoreceptors appeared disorganized and misshapen, contrasting with the uniform shape, radial orientation, and organization observed in the outer segments of photoreceptors in wild-type control retinas. Photoreceptors in wild-type larvae exhibited well-developed, long, calyceal processes. Notably, we observed a reduction in intact calyceal processes surrounding the outer segments in *pcdh15b* mutants compared to their wild-type siblings, with the calyceal processes appearing branched and thick. Using an antibody against human PCDH15, we observed stippled labeling in the region of the calyceal processes in sectioned retinas from wild-type larvae (Fig. 2I). This labeling was undetectable in *pcdh15b* mutant retinas (Fig. 2J). The morphological disruption suggested that photoreceptor function might be compromised, so we tested visual function using the optokinetic response assay in 5 dpf *pcdh15b* mutant larvae. Compared to wild-type controls, mutants consistently tracked fewer targets (Fig. 2K).

**Figure 2.**
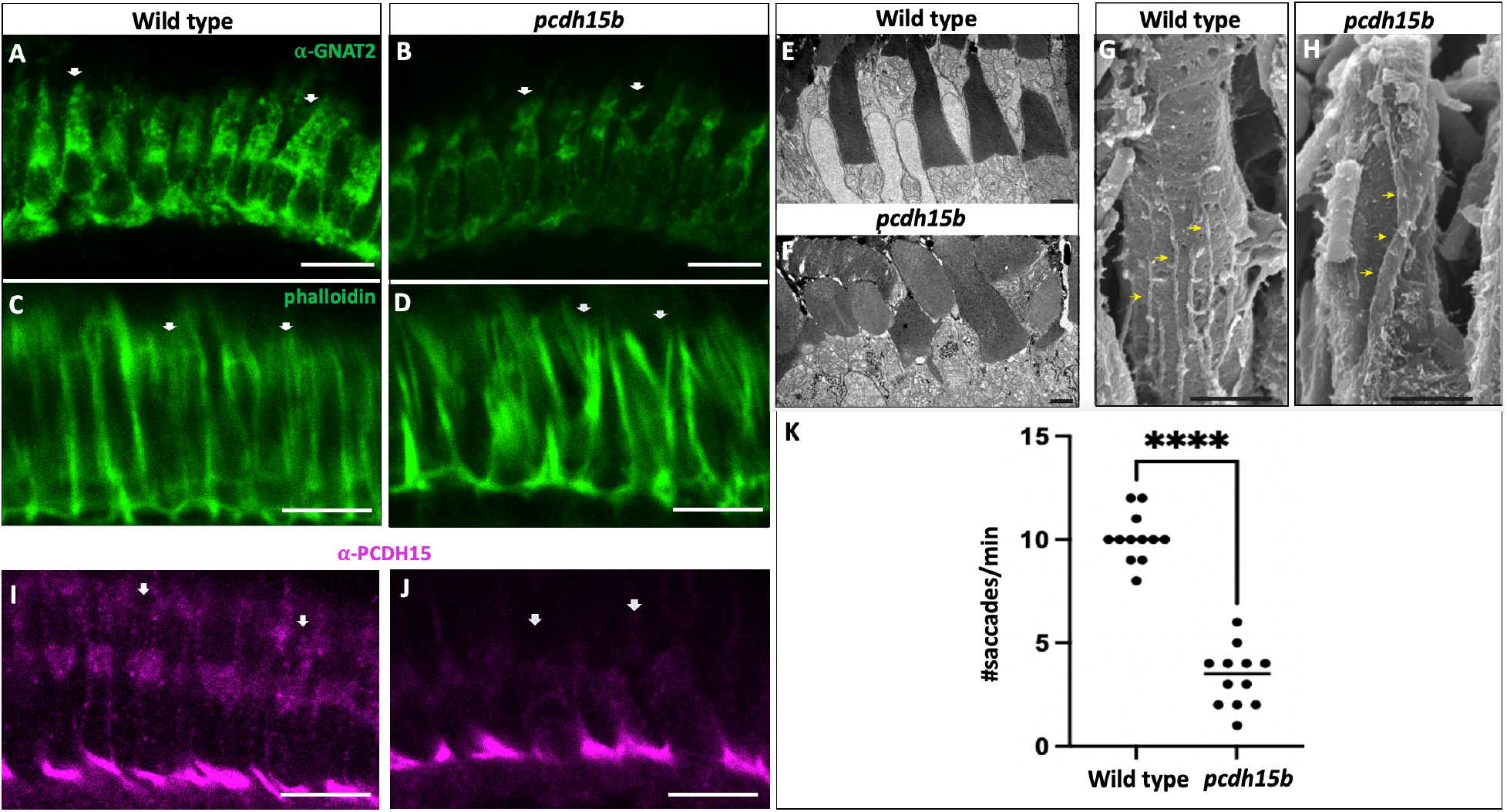
pcdh15b mutants have disorganized photoreceptors and reduced visual function. (A) Cone transducin labeled by the GNAT2 antibody fills the outer segments of wild-type cones (arrows) and outlines the cell body and pedicle in 6 dpf larval photoreceptors. (B) Cone transducin signal is diminished in *pcdh15b* mutant photoreceptors, revealing twisted and foreshortened outer segments. (C) Phalloidin labels actin filaments that provide structure to the photoreceptor outer segments, including the calyceal processes (arrows) arranged in an orderly fringe around the circumference of the cells. (D) Phalloidin-labeled calyceal processes are disorganized, fused, and splayed around the outer segments of *pcdh15b* mutant photoreceptors. (E-H) Representative transmission electron micrographs and scanning electron micrographs. In *pcdh15b* larvae, photoreceptor outer segments exhibit disorganized and misshapen morphology (F), in contrast to the well-organized, uniformly shaped, and radially oriented outer segments in wild-type retinas (E). Control photoreceptors have elongated calyceal processes (yellow arrows, G), whereas *pcdh15b* mutants show a notable reduction in intact calyceal processes surrounding the outer segments, and the few that form display an abnormal branched and thickened morphology (yellow arrows, H) compared to wild-type processes (G). (I) An antibody against human PCDH15 labels 6 dpf larval zebrafish photoreceptors at the base of the outer segment (arrows), corresponding to the region of origin of the calyceal processes. (J) PCDH15 antibody signal is undetectable at the region of the calyceal processes in *pcdh15b* mutant photoreceptors. Signal persists at the synapse, likely due to protein isoforms unaffected by this mutation. (K) Optokinetic response assay. An average of 10 saccades are recorded from 5 dpf wild-type larvae viewing stripes inside a rotating drum for one minute at 5 rpm. *pcdh15b* mutants track fewer targets (median 3.5) and show a larger variability in this response (*n* = 6 fish, tested in both directions). Each dot represents the number of saccades recorded from a single animal. *P* <0.0001. Scale bars: A-D 5 µm; E-F 2 µm; G-H 1 µm; I-J 5 µm.

We also analyzed photoreceptor cell death in 7 dpf larvae with anti-active caspase 3 antibody labeling to detect dying cells^[9]^. Caspase-positive photoreceptors were present in *pcdh15b* mutant retinas at more than five times the level observed in wild-type controls (Fig. 3).

**Figure 3.**
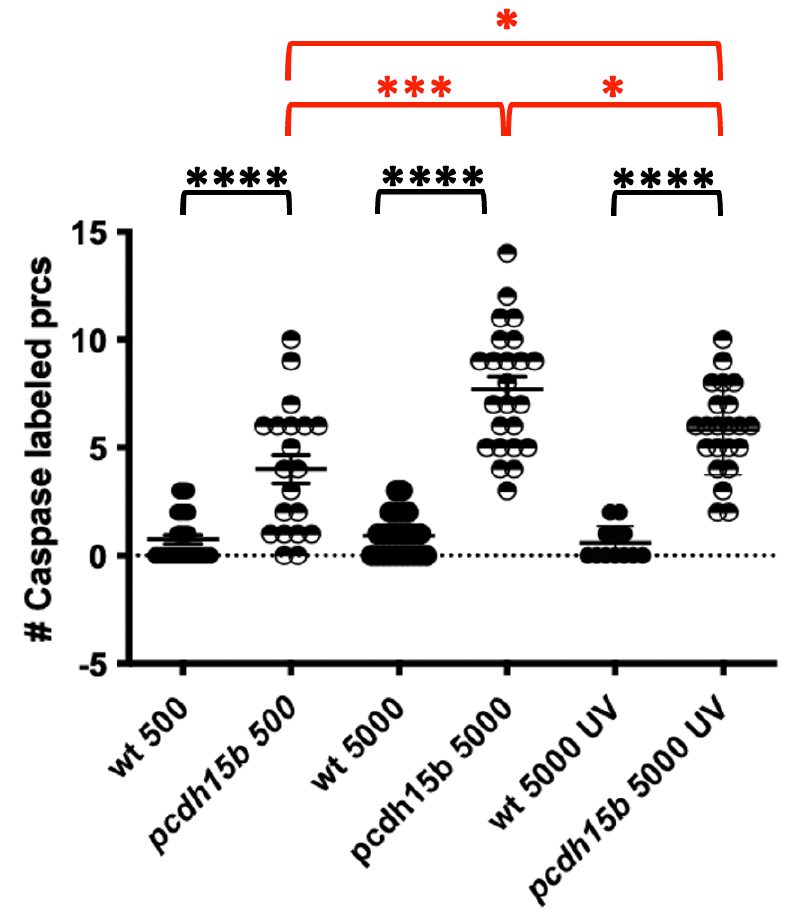
Bright light exacerbates photoreceptor cell death and a UV filter protects photoreceptors from cell death in *pcdh15b* mutants. Each dot represents the number of photoreceptor cells counted in serial sections of wild-type (filled circles) and mutant (half-filled circles) 7 dpf larval retinas after exposure to various light conditions. Wild-type retinas have a low (<1 cell per eye on average) level of cell death whether in normal facility conditions (500 lux), elevated intensity (5000 lux), or 5000 lux plus UV filter. In contrast, *pcdh15b* mutants show elevated retinal cell death compared to wild type even in facility light. This phenotype is enhanced under the higher light level, and attenuated by UV filter treatment. *n* ≥20 eyes for each group. Significance brackets between wild type and mutant are in black, between mutants under different conditions are in red. ***** *P* <0.0001; *** *P* = 0.0004; * *P* = 0.03 and 0.04.

### Exposure to higher light intensity exacerbates photoreceptor cell death in *pcdh15b* mutants

We previously demonstrated a relationship between photoreceptor death and light exposure in other zebrafish models of USH^[9]^, and raising USH mutants in the dark has been shown to be protective^[9, 20, 22, 31]^. We tested whether this phenomenon also occurs in our *pcdh15b* mutant lines. Mutants and wild-type controls were maintained in Petri dishes for 3 days starting at 4 dpf and exposed either to standard fish facility illumination of 500 lux or to elevated light exposure of 5000 lux using a broad spectrum bulb that extends to UVB wavelengths. All animals were maintained on a 14 hour light-10 hour dark cycle.

No increase in photoreceptor cell death was noted in wild-type larvae raised at the higher light intensity compared to wild types raised at normal light intensity (Fig. 3). However, *pcdh15b* mutants had increased cell death compared to wild-type controls raised at normal 500 lux facility light intensity. The amount of photoreceptor cell death in *pcdh15b* mutants was even greater at the higher 5000 lux intensity compared to mutants raised at the lower light intensity and compared to wild-type controls at both intensities. To explore the contribution of high energy wavelengths to this light-induced photoreceptor degeneration in *pcdh15b* mutants, we raised a subset of animals in Petri dishes with lids covered by UV-blocking window film. When exposed to the same higher 5000 lux light intensity of the broad spectrum bulb, which once again had no effect on the low level of caspase 3 activity in wild-type retinas, mutant larvae receiving UV protection had reduced photoreceptor cell death compared to unprotected mutants (Fig. 3). Together these results show that photoreceptor cell death in *pcdh15b* mutants is exacerbated by bright light exposure, and blocking UV can significantly protect the mutant retina.

Our finding of elevated photoreceptor cell death on exposure to higher intensity light (Fig. 3) suggested that cell stress may play a role in the degenerative process of our USH1F fish model. Evidence of ocular oxidative damage and ongoing oxidative stress in Retinitis Pigmentosa patients has been clinically documented^[7]^. Moreover, the combination of endogenous and exogenous oxidative stress produced during normal retinal metabolism and the mounting metabolic challenges of degenerative retinal diseases can be both causative and consequential factors in the disease pathology^[8]^. Our USH1F model, with a strong retinal degeneration phenotype that can be modulated by light exposure, provides a platform to test the effects of antioxidant treatment on this pathophysiological process.

We focused our studies on hexafluoro^[5]^, a synthetic biphenolic compound we developed^[5]^ that promotes neuronal survival and differentiation by reducing oxidative stress through activation of the SIRT3 pathway^[1, 26]^. We first titrated the maximum tolerated dose by adding a range of hexafluoro concentrations, suspended in DMSO, to the growth medium of young wild-type fish beginning at 2 dpf. The medium was replaced daily through 6 dpf with new doses of hexafluoro, and animals were monitored for viability and developmental milestones each day. An equal number of control animals was raised alongside these experimental groups and exposed to the same concentration of DMSO without hexafluoro added to the growth medium. The maximum dose at which no developmental or behavioral abnormalities were observed during the testing period was 0.125 µM. This dose was then applied to mixed progeny from a cross of *pcdh15b*^*+/-*^ heterozygous carriers for the four-day testing period. At 5 dpf, the timepoint at which homozygous mutants can be distinguished from heterozygotes and homozygous wild types by their abnormal swimming behavior, animals were washed to remove the drug and assayed for optokinetic response.

Like untreated mutants (Fig. 2K), mutants treated with DMSO alone showed diminished visual responses (Fig. 4A). Mutants exposed to hexafluoro displayed a significant improvement in visual function, which, although still below wild-type responses, was nevertheless notably increased compared to mutants from the DMSO control group. To measure the loss of photoreceptors (including non-caspase mediated cell death), we cut thin plastic sections of larval retinas from the various experimental and control groups, stained them with Toluidine Blue, and counted photoreceptor nuclei in a consistent region the central retina (Fig. 4B). We measured an increase (*P* = 0.0128) in the number of cells retained in treated versus untreated mutants (Fig. 4C). However, because the drug testing during this experiment was carried out at extremely low light levels (30 lux), we reasoned that brighter light exposure might better reveal the efficacy of hexafluoro treatment. Our data implicating environmental light levels in the rate of photoreceptor cell death in *pcdh15b* mutants (Fig. 3) suggested the possibility that the low illumination in this experiment was protective, and the protective environment could underly the improved visual function recorded from the mutants. This protective effect might diminish or disappear with the increased retinal cell stress of an elevated light environment.

**Figure 4.**
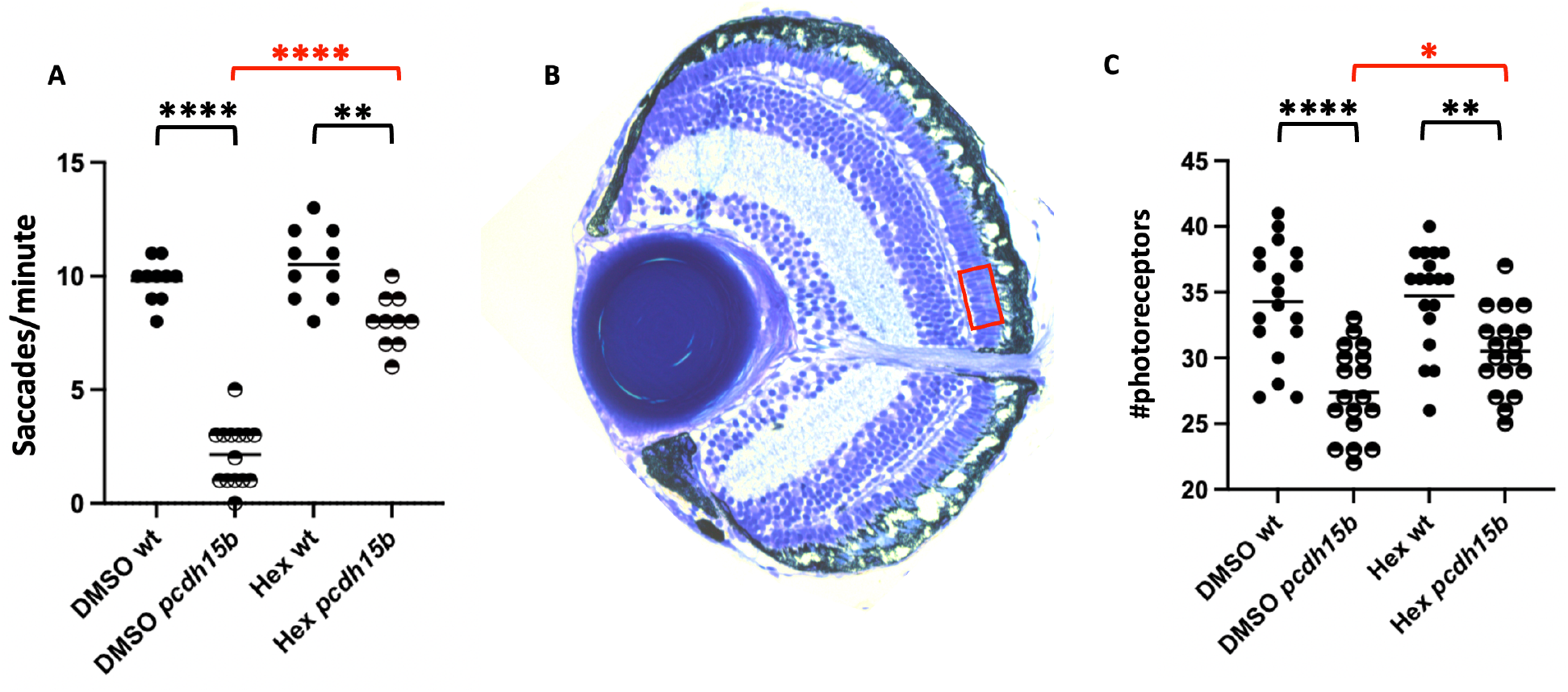
Hexafluoro improves visual function and photoreceptor survival in *pcdh15b* mutants. (A) Visual function. Each dot represents the number of saccades per minute recorded from a single 5 dpf larva (*n* ≥5 for all groups) tested with clockwise and counterclockwise rotation, producing two data points per individual fish. Saccades recorded from wild types (filled circles) and mutants (half-filled circles) treated with DMSO are consistent with previously reported data (Fig. 2G). Saccades recorded from mutants treated with hexafluoro are significantly elevated compared to untreated mutants. (B) Image of a toluidine blue stained 2 µm plastic section of a medial slice through a 7 dpf larval retina. The 50 µm x 20 µm standardized area outlined in red delineates the area superficial to the optic nerve in which cells were counted. (C) Photoreceptor survival. Each dot represents the total number of cells counted in the standardized area from each eye (*n* = 18 for each group, two eyes from each of 9 fish). Significance brackets between wild-type and mutant groups are in black, significance brackets between treated vs. untreated mutants are in red. A: **** *P* <0.0001; ** *P* = 0.002. C: **** *P* <0.0001; ** *P* = 0.001; * *P* = 0.01.

To test the range of hexafluoro’s effects on visual function and on the potential protective role of a low light environment, and to determine whether there is a correlation between improved function and the number of surviving photoreceptors, we raised groups of larvae either in standard facility lighting of 500 lux, or the elevated light level of 5000 lux (Fig. 5). As in the original experiment (Fig. 4A), mutants exposed to hexafluoro in DMSO performed better in the optokinetic response assay compared to mutants who received DMSO only, regardless of the intensity of light exposure (Fig. 5A, C). Additionally, we detected a significant increase in the number of photoreceptor cells retained in mutants exposed to hexafluoro raised at both higher light levels (Fig. 5B, D). Together these results show that exposure to brighter light exacerbates photoreceptor cell death and loss of visual function in *pcdh15b* mutants, and that hexafluoro treatment significantly protects the retina from these effects.

**Figure 5.**
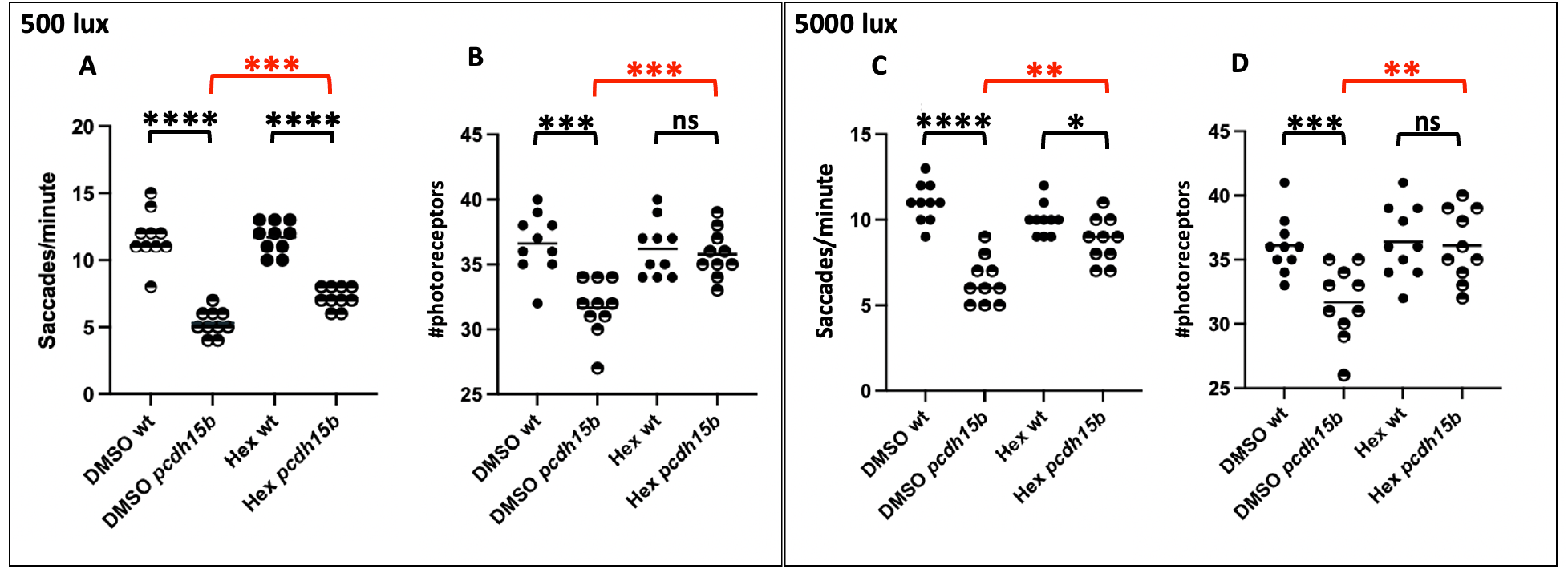
Hexafluoro rescues photoreceptors in *pcdh15b* mutants. (A, C) Optokinetic response assay results from 5 dpf larvae raised in standard facility lighting (A) or at the 5000 lux elevated light level (C). Each circle represents a response from an individual fish tested in clockwise and counterclockwise rotations (*n* = 5 fish per group). (B, D) Photoreceptor cell counts from 7 dpf wild-type and mutant animals reared at facility (B) or elevated (D) light conditions. Each circle represents the number of photoreceptor cells counted within the standardized area (Fig. 3B) of a sectioned retina (*n* = 10 eyes from 5 fish). Data points from wild-type fish in each graph are represented by filled circles, data points from mutants are represented by half-filled circles. Significance: brackets between wild -ype and mutant groups are in black, brackets between treated vs. untreated mutants are in red. A: **** *P* <0.0001; *** *P* = 0.0006. B: *** *P* = 0.0001. C: **** *P* <0.0001; ** *P* = 0.002; * *P* = 0.05. D: *** *P* = 0.0009; ** *P* = 0.004.

## DISCUSSION

We describe a comprehensive model of USH1F with both inner ear and retinal dysfunction, and have documented defects in hair cell morphology and vestibular function not previously seen in either morpholino^[28]^ or mutant^[20]^ studies. The combined defects in hair cells and photoreceptors documented in these *pcdh15b* mutant animals present the most complete model of USH1F to date. This model provides a unique opportunity to explore underlying pathophysiological mechanisms of the disease, as well as an excellent platform to test therapeutic intervention.

The reasons for the more severe and complete phenotypes in our mutants is not entirely clear. One difference is the position of the genetic lesion. It may be that our alleles, designed specifically to model the R245X mutation in human exon 8, alter a transcript expressed in hair cells that was left intact in other models. Although little is known about the number or variety of zebrafish *pcdh15b* transcripts, the human *PCDH15* gene produces dozens of splice variants. The exon structure of both zebrafish *pcdh15* genes is extremely well conserved compared to the human gene, indicating the possibility that multiple splice variants are also generated in fish. It is also possible that the mutations generated in the previous study^[20]^ resulted in genetic compensation^[11]^. Despite the differing effects on balance, the results of all of the morpholino and genetic purturbations are consistent with the interpretation that development and maintenance of normal hearing in zebrafish require *pcdh15a* function. However, we cannot rule out a supporting role for *pcdh15b* in hearing. The utricular hair cells are reported to respond to lower frequencies compared to those of the saccular hair cells, and the combined responses of cells in both areas may also contribute to acoustic sensation^[29]^. The stereocilia-specific defects we report here could reasonably impact perception of frequency range and amplitude.

Our study also expands our previous body of work describing the impact of environmental light conditions on retinal degeneration in zebrafish USH models^[9, 10, 24]^. Although some zebrafish models require elevated light rearing to exhibit a retinal degeneration phenotype, our *pcdh15b* mutants have a more severe yet still light-modulated retinal phenotype. In the *pcdh15b* mutants there is notable photoreceptor cell degeneration at ambient indoor light conditions, and cell death is significantly exacerbated by exposure to moderate light levels (5000 lux) equivalent to outdoor light on an overcast day. We also found that cell death is attenuated by the application of a UV filter. Although the precise mechanism for this dynamic effect is still being explored, the robust reproducibility of this phenomenon may have important near-term implications for USH patients, suggesting that judicious use of good-quality sunglasses is warranted.

Our findings that hexafluoro treatment can enhance visual function in USH models, regardless of the light environment in which the fish is raised, is extremely promising. Importantly, we show that USH1F fish models raised in ambient daylight levels, which approximate the diurnal living conditions of humans, show increased retinal cell retention after hexafluoro treatment. Unsurprisingly, hexafluoro does not improve the morphological defects observed in *pcdh15b* mutant photoreceptors even though the function and survival of these cells are significantly improved. Although the mechanism of this action of hexafluoro is still being investigated, the measurable and reproducible improvements observed in treated USH1F zebrafish models indicate that dysfunction downstream of the primary structural defects is at least in part regulated by oxidative stress. Previous studies of enteric neurons^[1]^, showed that hexafluoro treatment increased levels of neuronal nitric oxide synthase and choline acetyltransferase mRNA. Hexafluoro also significantly increased SIRT3 mRNA levels, and SIRT3 knock-down prevented the hexafluoro-mediated suppression of mitochondrial superoxide release. It remains to be shown whether hexafluoro acts through the same pathways to save photoreceptor cells in the USH mutant retina.

We suggest that hexafluoro and other compounds that promote cellular homeostasis by reducing reactive oxygen species could maximize function and survival of diseased cells. This could be significant for USH patients who are waiting for emerging targeted therapies that are still in preclinical development. Such treatments could also play an important clinical role when administered alongside cell-specific or gene-specific therapies targeting degenerative disorders.

## METHODS

### Animal husbandry

All live zebrafish experiments were performed using protocols approved by the University of Oregon’s Institutional Animal Care and Use Committee. Embryos and larvae housed in Petri dishes containing Embryo Medium^[34]^ were maintained at 28.5°C in the University of Oregon Zebrafish Facility on a 14 hour light/10 hour dark cycle. Embryos produced for these experiments were obtained through natural crosses of adult carriers.

### Generation of pcdh15b mutants

We designed and synthesized a gRNA, *pcdh15b* CRISPR 731F (aattaatacgactcactataGAGCTCCAAACCCGAAGGACgttttagagctagaaatagcaagttaaaataaggctagtccgttatcaacttgaaaaagtggcaccgagtcggtgcggatc) (MEGAscript® T7 Kit, Invitrogen) targeted to exon 8 of *pcdh15b* and injected this in combination with zebrafish optimized Cas9 mRNA synthesized from pT3TS-nCas9n^[15]^ into one cell-stage embryos spawned from wild-type Oregon ABC fish. At 24 hours postfertilization (hpf), 6-8 injected embryos were collected and pooled for DNA extraction. The resulting DNA templated a PCR reaction with primers flanking the region of desired CRISPR activity (pcdh15b e8-77f TTGGCTGAAGTGTATGTTTTGGAG and pcdh15b e8+38R GACAGAGCTGGGAAAGTAGGC) amplifying a 286 bp fragment. Amplicons were sequenced (Azenta Life Sciences) and traces were analyzed for the presence of disrupted reads at the anticipated cut site. Clutches from which positive samples were obtained were raised to adulthood at which time individuals were screened using DNA obtained from a fin clip using the genotyping procedure described above. Genetically confirmed carriers were bred to produce mutant offspring for analysis.

### Startle response assay

Free swimming 5 dpf zebrafish larvae in a 100 mm x 15 mm Petri dish filled with EM were observed from above. The startle stimulus was a brief series of sharp taps on the edge of the dish, as previously described^[21]^. Larvae responding with impaired swimming behavior were removed from the dish and subsequently genotyped to confirm homozygosity for the *pcdh15b* mutation. Abnormal swimming and impaired righting invariably correlated with a homozygous mutant genotype in repeated experiments (*n* >200). Swimming behavior was subsequently used to identify mutants for additional experiments.

### Optokinetic response assay

The assay was performed as previously described^[25]^ with the modification of extending the testing time to 1 minute in each direction.

### Hexafluoro treatment

Hexafluoro was dissolved in DMSO to 1 mM concentration and stored at -20°C. To identify the maximum tolerated dose for young zebrafish, stock solution was added to 5 ml EM per well in a 6-well plate, each containing eight 2 dpf wild-type larvae. Final concentrations ranged between 2 µM and 50 µM. Embryos or larvae were observed daily for viability and developmental abnormalities. A sharp titration curve was observed with these concentrations. Doses between 50 µM and 10 µM were 100% lethal within 24 hours, whereas larvae treated with 2 µM survived to 5 dpf, but showed signs of abnormal development including neurodegeneration, craniofacial abnormalities, and pericardial edema. In a second round of titrations ranging from 0.125 µM to 1 µM, a dose-response curve was again observed. At the highest dose, the same developmental disruptions were observed, but with a progressively later onset. At between 0.75 µM and 0.25 µM, larval morphology appeared normal but animals failed to inflate their swim bladders and exhibited impaired orientation. Animals treated with 0.125 µM showed no sign of disrupted development or behavior. The established experimental dose of 0.125 µM was then applied to 48 hpf embryos from *pcdh15b* mutant carriers in standard 100 mm Petri dishes with a total volume of 60 ml, replaced daily. Control dishes with the same number of animals and containing a volume of DMSO equal to the volume of Hexafluoro plus DMSO added to EM in the experimental groups were maintained alongside the hexafluoro treated dishes for the duration of the experiments.

### Elevated light exposure

Increased environmental light was produced by a 15 Watt T8 UVB terrarium lamps (Zoo Med) suspended over the shelf holding embryos or larvae in Petri dishes. The light fixtures were plugged into an outlet connected to the timing system governing the 14 h light/10 h dark cycle throughout the facility. Light intensity was measured with a light meter (Sper Scientific) at the beginning of each experiment and confirmed daily concurrent with cleaning and media changes. For UV film treatment, pieces of Gila glare control window film (Eastman) were cut to fit the lid of a standard 100 mm Petri dish and affixed with tension to avoid bubbles.

### Fluorescent labeling and immunohistochemistry on fixed tissue

For whole-mount phalloidin labeling, 5 dpf animals were collected into 1.7 ml microcentrifuge tubes, euthanized, and fixed in 4% PFA diluted in PBS-T for 2 hours at room temperature or overnight at 4°C. After 3x10 minute washes in PBS-T to remove fixative, larvae were incubated in 0.2% Triton-X 100 diluted in PBS overnight at 4°C to dissolve otoliths. Tissue was then washed 3x10 minutes in PBS-T and blocked in a solution containing 5% NGS and 1% BSA diluted in PBS-T for a minimum of 30 minutes and up to 2 hours at room temperature. Tissue was then incubated in Alexa-fluor conjugated phalloidin (Thermo Fisher) diluted 1:20 in block overnight at 4°C. After washing 3x10 minutes in PBS-T, the final wash was removed and replaced with Vectashield (Vector labs). Processed larvae were mounted in a lateral orientation, coverslipped, and imaged on a Leica SP8 Confocal Microscope. For antibody labeling, larvae were processed as previously described^[9]^ and incubated in primary and secondary antibodies as follows: phalloidin-488 or phalloidin-633 1:100 (Thermo Fisher); αGNAT2: 1:100 (Abclonal); α-active Caspase-3: 1:500 (BD Pharmingen); α-PCDH15: 1:200 (NSJ Bioreagents); Alexafluor Goat anti-rabbit 488 or 568 (Thermo Fisher).

### Toluidine blue staining of semithin plastic sections

Larvae were collected and fixed as described above, then embedded in acrylic resin (EMS) and cut into 2 µm sections that were collected serially on glass slides. Sectioned tissue was then covered with toluidine blue (Sigma Aldrich) dissolved in water and heated gently until the solution began to dry around the edges of the droplet. Slides were then rinsed 3-4 times in distilled water before being dehydrated, mounted in Xylene, and coverslipped. Images of stained retinas were obtained on an Aperio VERSA200 imaging system or Keyence BZ-X810 microscope and converted to image files for analysis.

### Transmission electron microscopy

At 7 dpf, larvae were fixed with 2.5% glutaraldehyde in 0.1 M cacodylate buffer (pH 7.2) for 24 hours at 4°C. After a triple rinse in cacodylate buffer, samples underwent postfixation with 1% osmium tetroxide/1.5% potassium ferrocyanide in 0.1 M cacodylate buffer for 24 hours at room temperature in the dark. Subsequently, samples were washed thrice in 0.1 M cacodylate buffer (pH 7.2), briefly rinsed in distilled water, dehydrated in an ascending series of ethanol concentrations, equilibrated in propylene oxide, and finally infiltrated and embedded in epoxy resin (Araldite 502/Embed-812 embedding media). Larvae were positioned in molds and resin polymerized at 60°C for 48 hours. The resin blocks were sectioned at 60–80 nm intervals using a Reichert Ultracut S ultramicrotome. Sections were mounted on copper Formvar/Carbon-coated grids and examined with a JEOL 1200EX microscope operating at 80 kV.

### Scanning electron microscopy

The retinal pigment epithelium and neural retina from enucleated eyes were quickly dissected apart and fixed with 2.5% glutaraldehyde in 0.1 M cacodylate buffer (pH 7.2) for 1-2 hours at room temperature. After a triple rinse in cacodylate buffer, samples underwent postfixation with 1% osmium tetroxide in 0.1 M cacodylate buffer for 4 hours at room temperature in the dark. After rinsing in 0.1 M cacodylate buffer (pH 7.2) and distilled water, the samples were dehydrated using an ascending ethanol series and critically dried with liquid CO_2_ (Tousimis Autosamdri 815). Subsequently, the samples were mounted on aluminum stubs with carbon conductive tabs, sputter-coated (Leica EM ACE600 sputter coater) with 5 nm platinum, and examined using a field-emission scanning electron microscope (Hitachi S-4700).

### Statistical analysis

Toluidine blue images opened in ImageJ were processed with the measuring tool to delineate the 50 µm x 20 µm area superior to the optic nerve, and a marking tool was used to mark and count individual photoreceptor nuclei within the selected area. Data collections were analyzed in Prism 10 and significance was determined through Mann-Whitney nonparametric tests.

## ACKNOWLDEGEMENTS

We gratefully acknowledge generous support from the Usher 1F Collaborative and the Usher Syndrome Society, and a Blavatnik Therapeutic Challenge Award..

